# Structural basis of how the BIRC6/SMAC complex regulates apoptosis and autophagy

**DOI:** 10.1101/2022.08.30.505823

**Authors:** Julian F. Ehrmann, Daniel B. Grabarczyk, Maria Heinke, Luiza Deszcz, Robert Kurzbauer, Otto Hudecz, Alexandra Shulkina, Rebeca Gogova, Anton Meinhart, Gijs A. Versteeg, Tim Clausen

**Affiliations:** Research Institute of Molecular Pathology, Vienna BioCenter, Vienna, Austria; Max Perutz Labs, University of Vienna, Vienna BioCenter, Vienna, Austria; Vienna BioCenter PhD Program, Doctoral School of the University of Vienna and Medical University of Vienna, Vienna BioCenter, Vienna, Austria

**Author notes:** Correspondence should be addressed to Julian F. Ehrmann; Daniel B. Grabarczyk and Tim Clausen. These authors contributed equally (Julian F. Ehrmann, Daniel B. Grabarczyk).

## Abstract

Inhibitor of apoptosis proteins (IAPs) bind to pro-apoptotic proteases, keeping them inactive and preventing cell death. BIRC6 is an exceptionally large, multidomain IAP that inhibits its targets by means of its atypical ubiquitin ligase activity and in addition, functions as an inhibitor of autophagy by depleting LC3B. Little is known of the mechanisms by which BIRC6 interacts with its targets and fulfills these two roles. Here, we determined the cryo-EM structure of BIRC6 alone and in complex with two mitochondrial pro-apoptotic proteins, HTRA2 and SMAC. We show BIRC6 is an antiparallel homodimer that forms a crescent shape that arcs around a spacious cavity. The cavity is surrounded by binding sites for client proteins, where they interact with the flexible UBC domain that mediates ubiquitin ligation. Functional data reveal that multivalent binding of SMAC in the central cavity obstructs substrate binding, impeding ubiquitination of both autophagy and apoptotic target proteins. Together our data reveal the molecular mechanisms of how SMAC specifically binds and inhibits BIRC6 to promote apoptosis, and how this regulatory mechanism also extends to autophagy substrates. The interaction sites are hot spots of cancer and atrophy mutations, highlighting the importance of carefully balancing the interplay between BIRC6 and SMAC.

---

Apoptosis is an evolutionarily conserved and essential form of programmed cell death used to remove damaged or surplus cells^1^. Dysregulation relates to disease, in particular cancer, atrophy, and neurodegenerative disorders^2,3^. To safeguard against untimely initiation of apoptosis, a family of functionally related proteins called the inhibitor of apoptosis proteins (IAPs) bind and inhibit caspases, cysteine proteases that execute the cell death program^4^. A common feature of all IAPs is the presence of a BIR domain, which binds to a specific N-terminal signal sequence, called an N-degron, in its target proteins^4^. Active caspases cleave themselves to expose internal N-degrons as the basis for IAP recognition. During apoptosis, damaged mitochondria elute the effector protein SMAC that also presents an N-degron and competes for BIR domain binding, thereby liberating caspases from IAPs^5,6^. Acting within the ubiquitin cascade of E1 activating, E2 conjugating and E3 ligating enzymes, most IAPs contain a RING domain that has E3 ubiquitin ligase activity^7^. This activity covalently modifies the target substrate with the small protein ubiquitin, directing it for degradation by the proteasome. Thus, binding of the IAP to its target both inhibits pro-apoptotic proteases and promotes their degradation. The IAP guardians themselves are tightly regulated, preventing premature responses to minor cell death stimuli^8,9^.

The human IAP BIRC6 (also known as BRUCE or APOLLON) is expressed in all tissues and is the only essential IAP family member^10,11^. Despite its giant size of 530 kDa, it has only two characterized domains, namely a BIR domain for N-degron binding and a UBC domain at its C-terminus, which is characteristic of E2 ubiquitin-conjugating enzymes. E2 enzymes commonly react with E3 ubiquitin ligases to transfer ubiquitin to the target protein^12^. The UBC domain of BIRC6 is homologous to those of UBE2O and UBE2Z, which function as hybrid E2–E3 enzymes mediating ubiquitin transfer without requiring a cognate E3 ligase^13-15^. Both the BIR domain and UBC domain of BIRC6 are important for its role in downregulating N-degron containing pro-apoptotic proteases such as caspases and HTRA2^16-19^. However, this regulation works in both directions, and these proteases also exploit the BIR domain to degrade BIRC6^16,17^. A distinguishing feature of BIRC6 compared to other IAPs are functions extending beyond apoptosis regulation^20-23^, exemplified by its ability to inhibit autophagy^20,24^ (**Fig. 1a**). During autophagy, ATG8 family proteins like LC3B recruit cargo carrying receptor proteins and form autophagosomal membranes when lipidated^25^. By ubiquitinating LC3B and causing its proteasomal degradation^24^, BIRC6 limits the availability of this important autophagic building block.

**Fig. 1.**
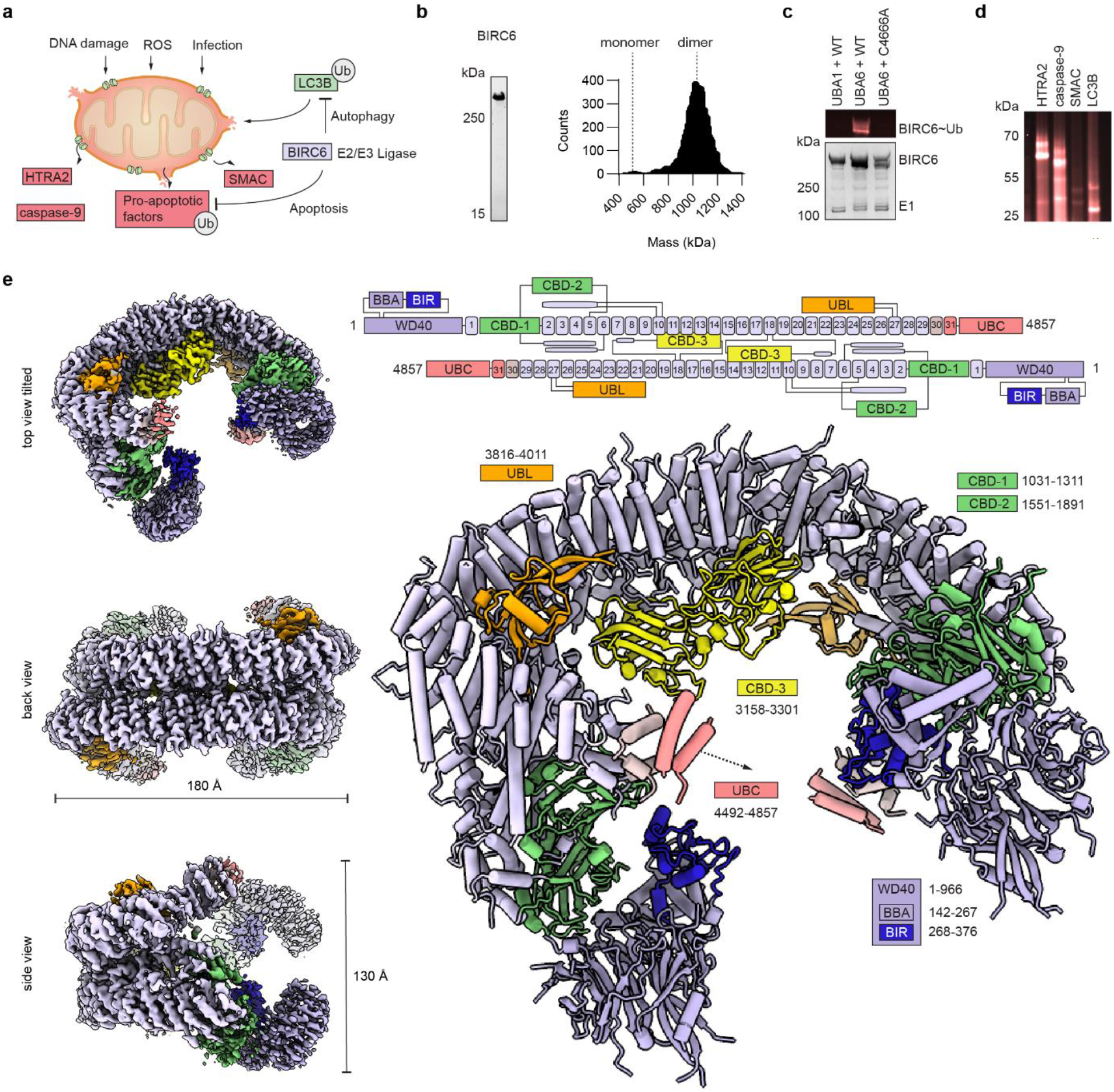
Cryo-EM structure of human BIRC6 in its active dimeric form. **a**, BIRC6 inhibits apoptosis and autophagy as an E2/E3 hybrid and ubiquitinated targets are marked for subsequent proteasomal degradation. **b**, Recombinant BIRC6 purity was assessed by SDS-PAGE, while mass photometry measurements at 20 nM revealed a dimeric species at ∼1.1 MDa. **c**, BIRC6 activity requires the cognate E1 UBA6, and is abolished when mutating the catalytic cysteine within the BIRC6 UBC domain (C4666A). DyLight488 labeled ubiquitin is spiked into the reaction for in-gel visualization (top). **d**, Ubiquitination assay of known interaction partners shows strong activity against HTRA2, caspase-9, and LC3B, but only low levels of SMAC ubiquitination. **e**, Cryo-EM map of BIRC6 in three orientations with dimensions indicated (left) and the model in cartoon representation (right). Domains are annotated with respective sequence boundaries and a schematic of the dimeric assembly is shown (top).

Currently, we have only a limited understanding of how BIRC6 targets and ubiquitinates diverse client proteins and how its IAP activity is regulated. Indeed, we have no structural data for any full-length IAP alone or in complex with their cognate substrates or regulators that would illustrate their molecular mechanisms. Whereas RING-containing IAPs represent dynamic particles that adopt diverse conformations and oligomeric states, in vitro characterization of BIRC6 is hindered by the enormous size of the protein. To address these knowledge gaps, we used cryo-EM to determine the structure of BIRC6 alone and in complex with the pro-apoptotic protease HTRA2 and the inhibitor SMAC. The structures reveal a head-to-tail homodimer in the shape of an open ring that encloses a central cavity used to accommodate clients. We show in molecular detail how HTRA2 and SMAC are captured in this central cavity, and how multivalent SMAC binding is used to regulate the specificity and activity of BIRC6. Furthermore, we show that LC3B binds to BIRC6 through sites distinct from apoptotic clients and is also subject to SMAC regulation. Together our findings demonstrate how the interplay between the non-canonical E2/E3 hybrid BIRC6 and the stress protein SMAC allows tight regulation of autophagy and apoptosis factors.

## BIRC6 dimerizes into a crescent scaffold with a central ubiquitination zone

To determine the architecture of BIRC6 and its molecular mechanism, we first expressed the human protein in insect cells. Mass photometry indicated that recombinant BIRC6 exists as a stable dimer, with a molecular weight of about 1.1 MDa (**Fig. 1b**). To test the activity of the dimer we performed ubiquitination assays in the presence of the obligate E1 ubiquitin-activating enzyme UBA6^16,24,26^ (**Fig. 1c**) and showed activity against reported substrates in vitro, validating the E2/E3 hybrid nature of BIRC6 (**Fig. 1a,d**). We then performed cryo-EM analysis of BIRC6 alone and in the presence of substrates. These data enabled us to generate a map of full-length BIRC6 with an overall resolution of 3.3 Å and build the structure of the active E2/E3 ligase (**Extended Data Figs. 1-5, Extended Data Table 1**). The structure revealed an intricate antiparallel dimer, whose four front-facing termini are connected by a twinned helical scaffold consisting of 31 armadillo repeats (**Fig. 1e, Extended Data Fig. 6a**). The resulting C-shaped structure has overall dimensions of 180×150×130 Å and encloses a central cavity. The N-terminal module is a seven-bladed beta propeller with two inserted adjacent domains (**Extended Data Fig. 6b,c**); a novel BIRC6 BIR-associated domain (BBA, 142-267), and the N-degron-binding BIR domain (268-376). The beta-propeller is flexibly linked to the start of the helical scaffold which is initiated by two carbohydrate binding-like domains, CBD-1 (1031-1311) and CBD-2 (1551-1891) (**Extended Data Fig. 6d**). Structurally related domains use a common surface patch to mediate interactions with carbohydrates or proteins, for example contributing to substrate recruitment in another ubiquitin ligase, the anaphase promoting complex^27^. Notably, for CBD-1 and CBD-2, these surfaces point to the interior of BIRC6 and may serve as accessory binding sites. A third CBD domain (CBD-3, 3158-301) is located at the center of BIRC6 where it homodimerizes via the described interface. Following this, a ubiquitin-like domain (UBL, 3816-4011) accompanies a sharp bend in the armadillo repeat backbone towards the front side, where the C-terminal UBC domain (4492-4857) is loosely tethered. Though we struggled to resolve the UBC domain in our EM reconstruction, in 2D classes and 3D variability analysis of a BIRC6 dataset in the absence of substrates we observed the appearance of globular density matching the dimensions of the UBC module extending from the C-terminal helical scaffold into the central cavity (**Extended data Fig. 6e**). The helical backbones of the two BIRC6 protomers interact intimately with each other, as reflected by the large interface of 9100 Å^2^ and the remarkable stability of the dimer (**Fig. 1b**). In addition to extensive van der Waals interactions between the stacked armadillo repeats, the dimer is stabilized by the interlocking of CBD-3 in the center of the protein. As well as forming a prominent protrusion in the central cavity, CBD-3 and an adjacent helix are swapped across the dimer, binding to the armadillo repeat of the other protomer (**Extended data Fig. 6f**). Overall, the dimer architecture is critical to orient the catalytic UBC and the substrate binding BIR domain towards the central cavity. This cavity has a diameter of about 50 Å, providing a spacious ubiquitination zone in reach of the mobile UBC heads.

## BIRC6 binds HTRA2 and LC3B through distinct sites

To visualize the substrate targeting mechanism, we reconstituted complexes with the client proteins caspase-9, HTRA2 and LC3B. Cryo-EM data of complexes with caspase-9 and LC3B were inconclusive with regard to substrate densities. We observed heterogenous densities in the central cavity which were similar to those detected in a BIRC6 dataset in the absence of substrates (**Extended data Fig. 7a**) and presumably originated from the flexible UBC domain or other low occupancy disordered regions. In strong contrast, the cryo-EM map of the stable complex of BIRC6 with HTRA2 revealed clearly defined features of the bound substrate (**Fig. 2a, Extended Data Fig. 7b-d**). The local resolution of HTRA2 was approximately 8 Å and due to its distinctive pyramidal shape the substrate could be unambiguously fit into the density (**Fig. 2a, Extended data Fig. 7c,d**). HTRA2 is a homotrimeric protein composed of a trypsin-like protease, a regulatory PDZ domain and one N-degron per protomer. Measuring about 66 Å per side, the trimeric protease core almost fully occupies the central cavity of BIRC6. The base of the trimer, where the proteolytic sites are located, packs against the two CBD-3 domains (**Fig. 2b, Extended Data Fig. 7e**). Both neighboring CBD-3s interact with HTRA2 using the same feature – an extended loop with three highly conserved arginine residues (**Fig. 2b**). These arginines recognize the charged surface area surrounding the active sites of HTRA2. In addition to undergoing electrostatic interactions with CBD-3, the protease is well positioned for its N-degrons to interact with the BIRC6 BIR domains, further anchoring the substrate to BIRC6. While we did not detect density for the N-degron, the six N-terminal residues are well positioned to bridge the 10 Å distance to reach the proximal BIR domain supporting a stable yet flexible binding mode.

**Fig. 2.**
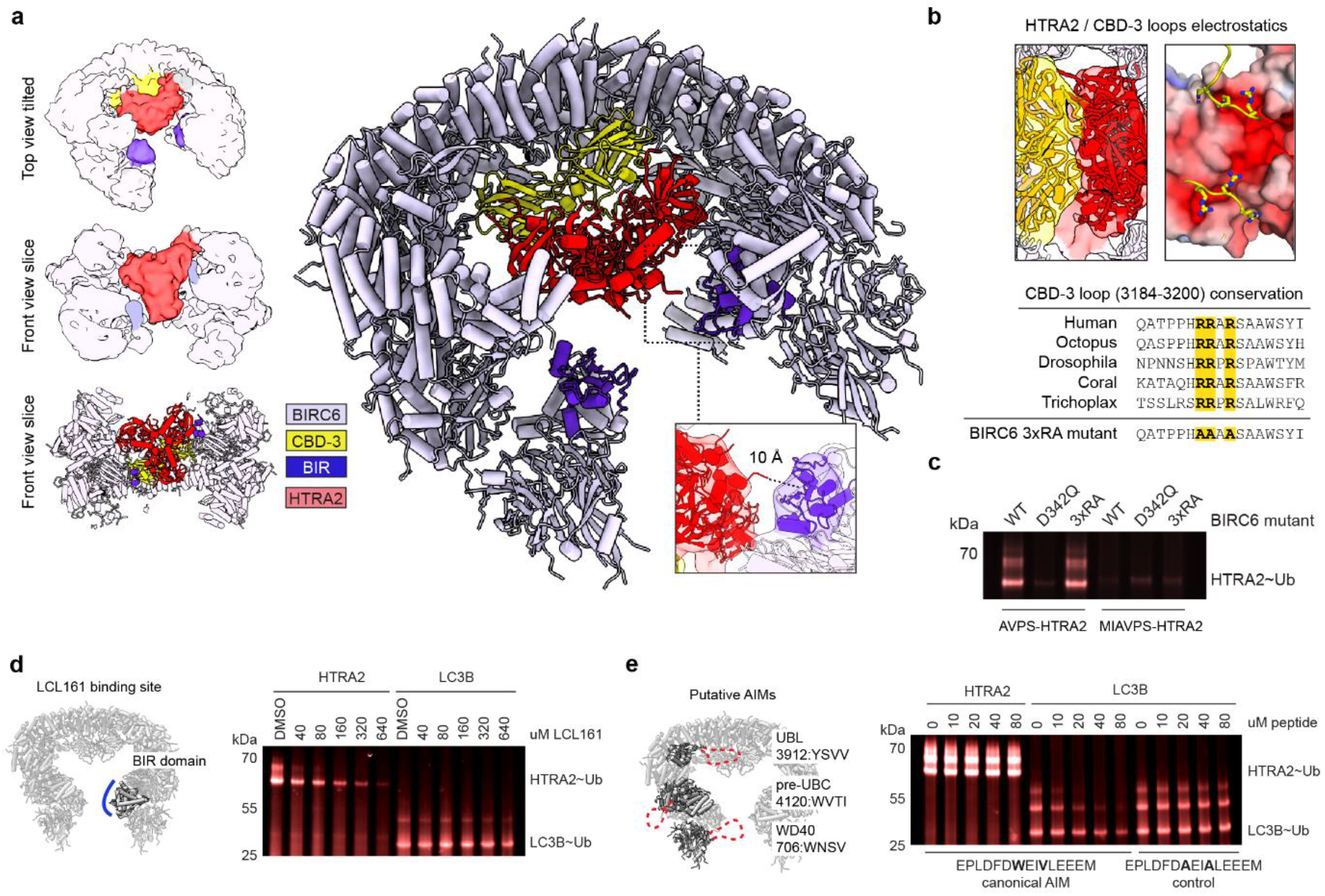
BIRC6 employs distinct targeting mechanisms for apoptosis and autophagy proteins. **a**, Cryo-EM structure of BIRC6 bound to HTRA2 lowpass filtered to 10 Å, color-coded as indicated and showing the characteristic trimer (left) with a cartoon representation highlighting the distance of the termini to the BIR domain (center). **b**, Cartoon and electrostatic representation of the CBD-3 arginine loops packing against the negatively charged protease. Representative alignment below highlighting the arginines, and the BIRC6 3xRA mutant (Arg3190Ala/Arg3191Ala/Arg3193Ala). **c**, Ubiquitination assay of HTRA2 with a native N-degron (AVPS) and a variant (MIAVPS) unable to bind the BIRC6 BIR domain. BIRC6-catalysed ubiquitination of HTRA2 mainly depends on the native degron, and the functional BIR domain. **d**, HTRA2 ubiquitination is sensitive to the small-molecule BIR domain competitor LCL161, which has no effect on LC3B modification. **e**, LC3B ubiquitination is decreased when titrating an AIM peptide into the reaction, while ubiquitination of HTRA2 was not affected.

To understand the contribution of the distinct interaction sites towards substrate binding and ubiquitination, we generated BIRC6 variants with either the N-degron binding site mutated (D342Q) or the CBD-3 arginine loop modified to negate the positive charge (3xRA). We observed that HTRA2 recognition was mainly dependent on the N-degron interaction, as D342Q abrogated ubiquitination (**Fig. 2c**). In agreement with this, adding the chemical N-degron mimetic LCL161 into the reaction as a competitor, or mutating the native N-degron of HTRA2 impaired ubiquitination. (**Fig. 2d**). Though not critical for ubiquitination, the interaction with CBD-3 could modulate HTRA2 function. In the previously described HTRA2 resting state, the PDZ domains are closely packed against the protease body blocking access to the proteolytic sites^28^. Moreover, NMR data indicated that binding of activator peptides leads to PDZ displacement and subsequent protease activation^29^. In our cryo-EM structure, the BIRC6 CBD-3 domains have displaced the PDZ domains, and density for one of these flexible domains can be detected outside of the trimer core (**Extended Data Fig. 7f**). Thus, binding to BIRC6 could be directly coupled with protease activation, providing a molecular mechanism of how HTRA2 selectively degrades IAPs upon mitochondrial stress^17^.

Considering its diverse cellular roles, we were curious how BIRC6 recognizes substrates outside the apoptosis pathway, especially those lacking an N-degron. The best characterized non-apoptotic target is the autophagy protein LC3B^24^. Though it has been shown that BIRC6 negatively regulates autophagy by ubiquitinating and thereby depleting LC3B, the targeting mechanism is unknown. One potential docking site is the BIR domain, which mediates diverse protein-protein interactions as exemplified by the phospho-dependent binding of histone 3 to BIRC5^30^. To test whether LC3B is also recognized by the BIR domain, we performed competition assays with the N-degron-mimetic LCL161 as well as using the D342Q variant of BIRC6. In contrast to HTRA2, these had no effect on LC3B ubiquitination (**Fig. 2d**). Since LC3B typically binds to proteins through ATG8-interacting motifs (AIMs)^22^, we performed a bioinformatic motif search and, after filtering for only those in disordered regions, identified several putative AIMs in BIRC6^31^ (**Fig. 2e**). We therefore asked whether targeting of LC3B may be mediated by such a motif and carried out competition experiments with a canonical AIM peptide. While the AIM peptide had no effect on targeting HTRA2, ubiquitination of LC3B was strongly reduced, suggesting that one or more of the putative BIRC6 AIMs recognizes LC3B (**Fig. 2e**). From these experiments, we conclude that BIRC6 has distinct targeting mechanisms for apoptotic and autophagy proteins.

## SMAC inhibits BIRC6 by multivalent binding to the central cavity

Although SMAC forms a stable complex with BIRC6 (**Extended data fig. 8a**), the apoptotic protein was only weakly ubiquitinated (**Fig. 1d**). We reasoned SMAC could compete, via its N-degron, with HTRA2 and caspase-9, and inhibit their ubiquitination. Indeed, we observed a complete loss of HTRA2 and caspase-9 ubiquitination even when these substrates were in 8-fold excess of SMAC (**Fig. 3a**). To our surprise, SMAC also completely inhibited the activity of BIRC6 against LC3B, despite the autophagy protein not binding to the BIR domain.

**Fig. 3.**
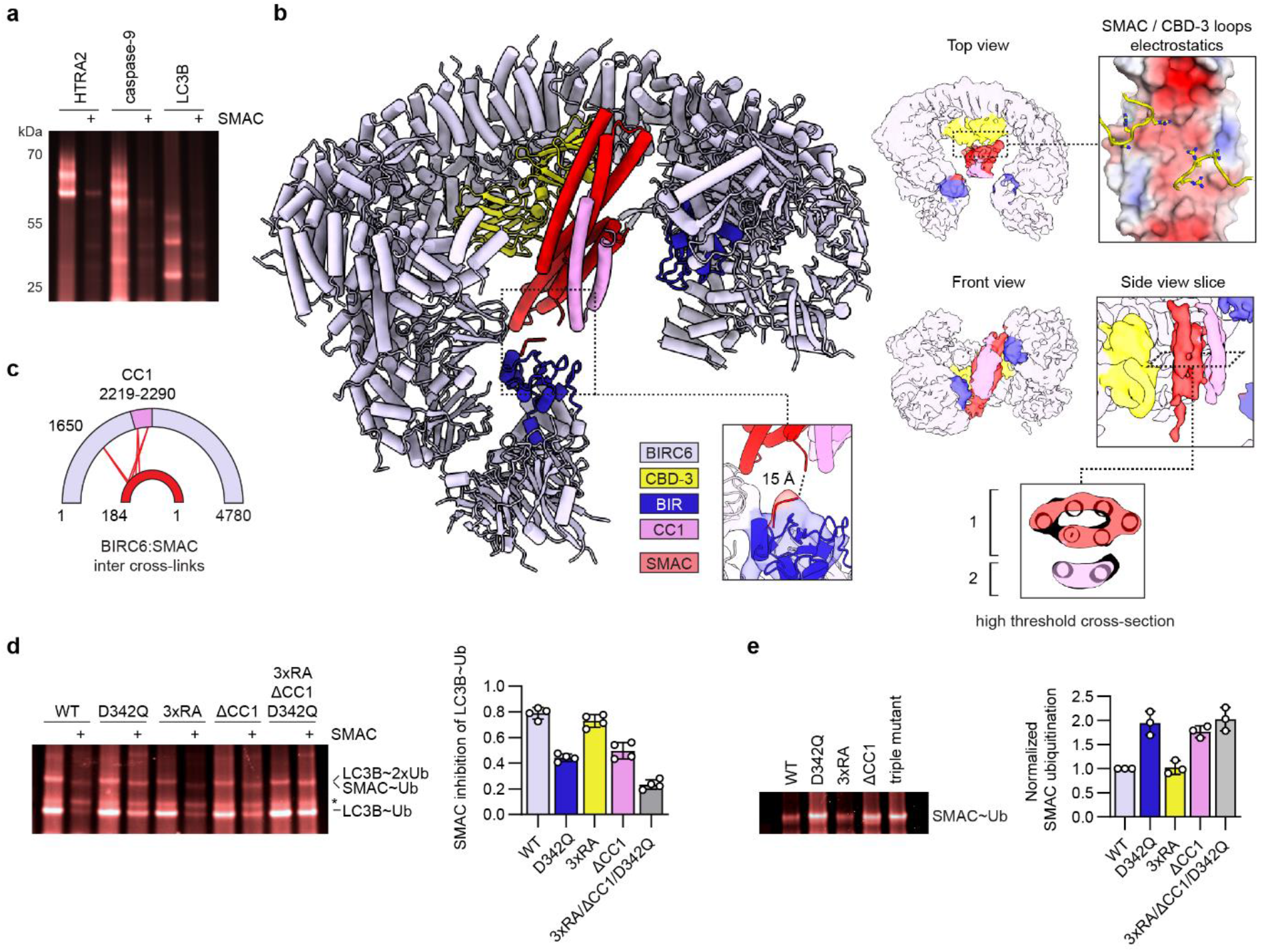
BIRC6 inhibition by SMAC is dependent on a specific tripartite binding mode. **a**, SMAC inhibits HTRA2, caspase-9 and LC3B ubiquitination. **b**, Cryo-EM structure of BIRC6 bound to SMAC with degron density highlighted. SMAC binds the CBD-3 loops with negatively charged surfaces reminiscent of HTRA2. The SMAC dimer (red) is wedged between the CBD-3 domains (yellow) and orphan density (purple). **c**, Cross-link coupled mass spectrometry detects multiple inter-molecular cross-links between SMAC and the CC1 insert (2219-2299). **d**, Using LC3B ubiquitination as functional readout, BIRC6 mutants with disrupted SMAC binding are less sensitive to SMAC inhibition. The fraction of the LC3B-Ub band lost upon SMAC addition is quantified for four replicates for each variant. **e**, SMAC is ubiquitinated more strongly by BIRC6 variants with disrupted interaction sites. Quantification of three replicates are plotted with SMAC ubiquitination normalised to the wild-type reaction.

To better understand the inhibitory mechanism of SMAC, we determined the structure of the BIRC6:SMAC complex by cryo-EM. SMAC was resolved at a local resolution of approximately 8 Å, enabling placement of dimeric SMAC into the density (**Fig. 3b**; **Extended data Figs. 8b**). Our structure shows that the six helices of the SMAC dimer penetrate the entire cavity of BIRC6, occluding the CBD-3 and BIR domains. Like HTRA2, SMAC is held in the center of the cavity by non-specific interactions of a negatively charged surface with the arginine loops of the two CBD-3 domains (**Fig. 3b**). Notably, this particular surface on SMAC has previously been reported to interact with another IAP, XIAP^32^, pointing to a shared binding mechanism to modulate IAP function (**Extended data Fig. 8c**). Both N-termini of the SMAC dimer protrude towards the BIR domains, and in this case low-resolution density for the bound N-degron can be observed (**Fig, 3b, Extended data Fig. 8d**). Importantly, an additional density corresponding to two helices was observed packing against the SMAC dimer (**Fig. 3b**). The extra helices could not be reconciled with the reported tetrameric form of SMAC^33^. We asked whether flexible segments of BIRC6 may contribute to this density, and employed cross-linking coupled mass spectrometry to identify the putative additional SMAC binding site. We detected 4 specific crosslinks between SMAC and a partially disordered insert in the helical backbone of BIRC6 that we termed coiled-coil 1 (CC1, residues 2219-2299, **Fig 3c**). This insert is predicted by Alphafold2 to form two antiparallel helices that fit well to our orphan EM density and has the right linker length to reach the modelled position (**Extended data Fig. 8e,f**). In conclusion, SMAC coordination relies on three main interactions: N-degron binding at the BIR domain, electrostatic interactions with CBD-3, and formation of a helical bundle with CC1.

LC3B is inhibited by SMAC despite not sharing any of the identified SMAC-BIRC6 interaction sites. To understand the mechanism of this inhibition, we mutated each interaction site and monitored the effect of SMAC on BIRC6-mediated LC3B ubiquitination (**Fig. 3d**). Disruption of the BIR domain with the D342Q mutation did not fully restore LC3B ubiquitination, suggesting the other two sites contribute to SMAC binding. The 3xRA mutation had little effect on SMAC inhibition, but deleting CC1 caused a moderate loss of inhibition, confirming the importance of this interface. A BIRC6 variant mutated in all three sites showed the strongest loss of SMAC mediated inhibition (**Fig 3d**). We conclude that multivalent interactions arrest SMAC in the central cavity, obstructing the ubiquitination zone and inhibiting BIRC6 activity.

As SMAC itself is poorly ubiquitinated, we tested whether its tripartite binding mode may be causative. We addressed this by ubiquitinating SMAC in the absence of other substrates using the same BIRC6 interface mutants (**Fig. 3e**). The D342Q mutation and the CC1 deletion resulted in a two-fold increase in SMAC ubiquitination, suggesting that the tight, multivalent tethering of SMAC also functions to protect itself from becoming ubiquitinated. We observed no further increase in ubiquitination for the variant mutated for all three sites, presumably due to overall weakening of the SMAC:BIRC6 interaction. In sum, these data suggest that the three characterized SMAC interaction sites are critical to limit its ubiquitination and allow for robust inhibition of BIRC6.

## Conclusion

IAPs function as cellular safeguards controlling programmed cell death. To achieve this task, IAPs must ensure that caspases are kept inactive under non stress conditions, while allowing their SMAC-induced release during apoptotic stimuli. To address the molecular mechanism underlying this fundamentally important decision on cell survival and cell death, we performed a mechanistic analysis of BIRC6, the only essential IAP. As revealed by its cryo-EM structure, the giant ubiquitin ligase adopts a C-shaped fold around a central cavity containing various receptor sites. The cavity fosters competition among substrates, with SMAC utilizing most of the provided contact points and thus outcompeting other clients. We thus propose that SMAC, upon release from damaged mitochondria, undergoes multivalent interactions with IAPs like BIRC6 to free the sequestered caspases, which then complete the cell death program. Moreover, multivalent binding protects SMAC from being ubiquitinated itself, explaining how the inhibitor can persist when bound to BIRC6 and ensure completion of apoptosis. On the contrary, under non-stressed conditions when SMAC is absent, BIRC6 can target and ubiquitinate caspase substrates. This basal activity is critical to prevent untimely apoptosis due to stochastic leaking of pro-apoptotic factors from mitochondria (**Fig. 4a**)^17^.

**Fig. 4.**
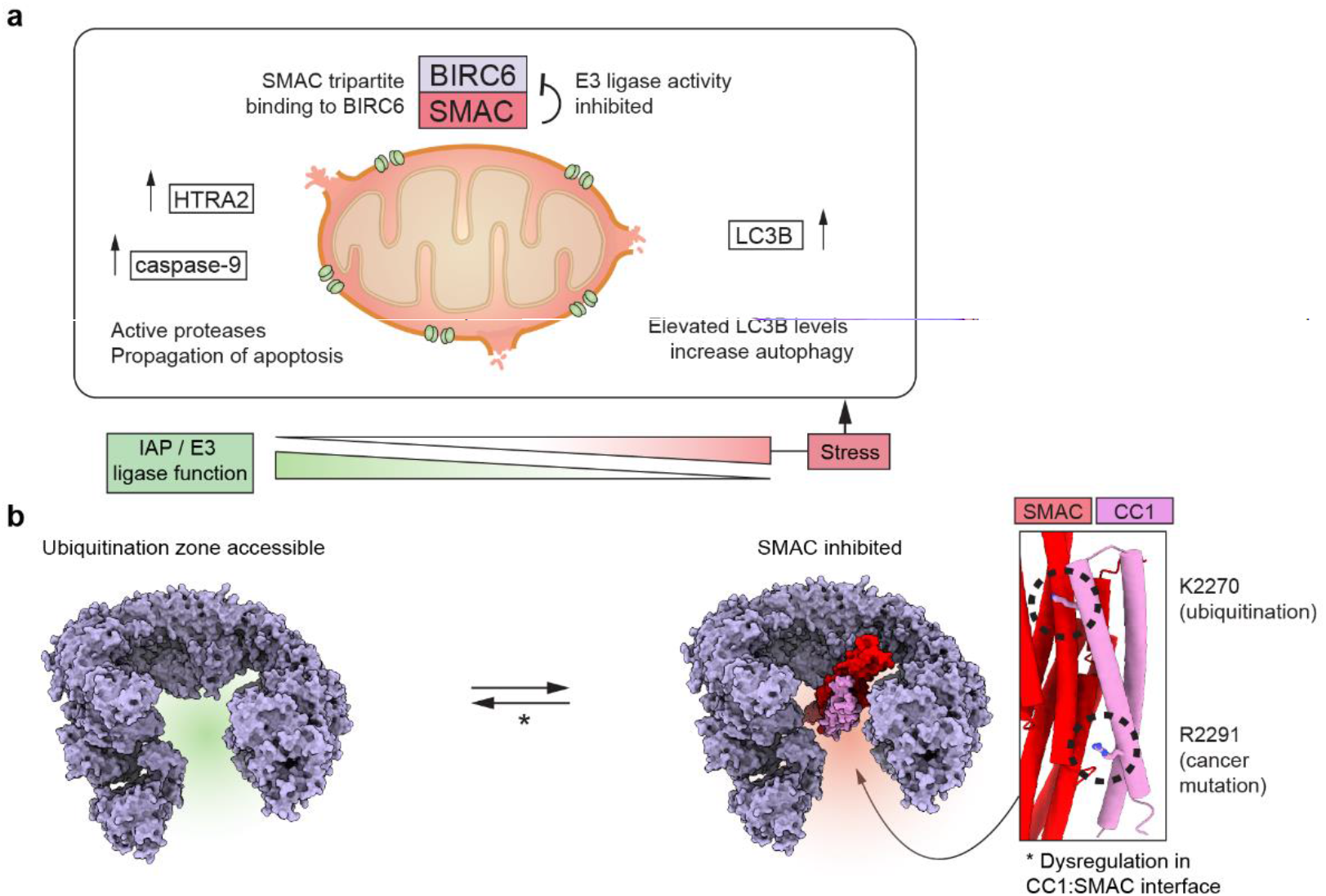
SMAC controls BIRC6 activity. **a**, During mitochondrial stress, cytosolic levels of SMAC increase, leading to inhibition of BIRC6 through multivalent binding. In this way, other N-degron substrates of BIRC6 like caspase-9 and HTRA2 are free to drive cell death, and LC3B can participate in autophagy. **b**, The BIRC6 CC1:SMAC helical bundle, which stabilizes the inhibited state, is prominently mutated in cancer and subject to post-translational modifications (inset), likely shifting the equilibrium to the active IAP.

The regulation and activity of BIRC6 is tied closely to key cellular processes. Clearance of toxic aggregates, for example, benefits from BIRC6 knockdown, as elevated ATG8 levels bolster the autophagy response. By studying the ubiquitination of LC3B, we show that BIRC6 uses AIM-like motifs to target ATG8 proteins. These motifs do not overlap with the binding epitopes of HTRA2 and other apoptotic proteases, allowing a separate regulation of cell death and cell survival pathways. Notably, SMAC inhibits ubiquitination of both apoptotic factors and LC3B. We reason that SMAC serves as a general regulator to switch BIRC6 activity on and off in response to stress stimuli requiring co-regulation of autophagy and apoptosis pathways^34,35^.

Our structural data pave the way for functional contextualization of known post-translational modifications and disease mutations within BIRC6 (**Fig. 4b**). For instance, an Arg3204Gln mutation in mice has been linked with developmental disorders^36^. The corresponding Arg3190 in humans is part of the CBD-3 arginine loop that interacts with the negatively charged surfaces of the HTRA2 substrate. Accordingly, the mutation may influence the substrate spectrum of BIRC6 or, alternatively, affect the binding mode and thus functional state of the pro-apoptotic protease HTRA2. Further, proteomics studies identified ubiquitination of Lys2270 within the SMAC interface of CC1 as the most abundant post-translational modification, possibly altering BIRC6 function^37^. A regulatory role of CC1 is consistent with our biochemical data showing that loss of CC1 correlates to decreased inhibition by SMAC. In line with this, CC1 residues in the SMAC interface are the most frequently substituted BIRC6 residues detected in cancer^38^. The most mutated site, Arg2291, is one of several conserved basic residues interacting with the negatively charged SMAC molecule. We hypothesize that CC1 mutations hinder SMAC binding and prevent apoptosis initiation. Tumors often present elevated levels of IAPs as a broad survival mechanism. Mutations disrupting the tripartite interface may thus desensitize cells to SMAC-triggered apoptosis, without impairing other crucial functions of BIRC6 for cell survival and tumor propagation. It appears that multivalent binding of SMAC offers an additional layer to regulate BIRC6 activity, and balance mitochondrial stress, apoptosis and autophagy.

## Acknowledgements

We thank the Vienna BioCenter Core Facilities for their support, especially Harald Kotisch and Tom Heuser (Microscopy), Jana Neuhold (Protein Technologies), and the Mass Spectrometry department. We are grateful for measurement time and support of the EM facility at the Institute of Science and Technology Austria (ISTA), especially V.-V. Hodirnau. Editing support was provided by Life Science Editors. We thank all members of the Clausen lab for discussions. We thank the Dagdas lab (Gregor Mendel Institut) for gifting the AIM peptides and LC3B plasmid. We thank the Ehrmann lab (University Duisburg-Essen) for gifting HTRA2 plasmids. This project received funding from a Marie Skłodowska-Curie grant (agreement no. 847548 to D.B.G.), an FFG Headquarter grant (no. 852936), a DocFunds grant (FWF no. DOC 112-B to J.F.E.) and the Austrian Science Fund (FWF, no. SFB F 79 to R.G.). The IMP is supported by Boehringer Ingelheim.

## Author contribution

J.F.E., M.H., L.D., R.K., A.S., R.G. prepared expression constructs and performed protein purifications. J.F.E. and D.B.G. performed biochemical experiments. J.F.E. prepared and screened cryo-EM specimen and performed initial processing. D.B.G. performed cryo-EM processing and model building. O.H. performed cross-link coupled mass spectrometry analysis. A.M. and G.V. helped analyze data. J.F.E. and D.B.G., designed experiments and T.C. outlined the study. J.F.E., D.B.G., and T.C. prepared the manuscript with input from all authors.

## METHODS

### Cloning, protein expression and purification

The BIRC6 gene sequence was codon optimized for insect cell expression, avoiding BsaI restriction sites, and split into 6 fragments of similar length. Fragment boundaries were flanked with BsaI sites to create 4 base pair overhangs as GoldenGate compatible quadruplets for scarless assembly. Genes were ordered as sequence verified fragments in vector backbones with kanamycin resistance. A polyhedrin promoter was cloned into a pUC19-based backbone flanked with BsaI sites producing a 3’ CCAT overhang and a 5’ ACCT overhang. The SV40 terminator was cloned into a backbone with a similar procedure, with BsaI-generated 3’ GTAT and 5’ TGAC overhangs. The promotor, gene, and terminator fragments were assembled by GoldenGate reaction^39^ into a pGBdest vector using gentamycin selection for correct assembly. Any subsequent point mutations or deletions were generated via Q5 mutagenesis within the single gene fragment spanning the region of interest as previously described^40^. Final constructs were then transformed into DH10EmBacY cells using Eugene HD transfection reagent (Promega) and recombinant bacmids were purified by isopropanol/alkaline lysis. Isolated DNA was then transfected into Sf9 cells (Expression Systems, 94-001) via Polyethylenimine PEI 25K (Polysciences, 23966-1) and V0 virus were subsequently amplified.

### Protein production in insect cells

BIRC6 wild type and mutants were produced with a C-terminal fusion to a Twin-Strep-tag. Baculovirus containing the desired constructs were supplied by the VBC Facilities. For expression, Trichoplusia ni High Five cells at a density of 1×10^6^ mL^-1^ in ESF921 serum-free growth medium were transfected with respective Baculovirus. Expression proceeded over three days at 27 degrees Celsius and 100 rpm, before harvesting the cells by pelleting at 500 g for 10 minutes and resuspension in 25 mM HEPES pH 7.5, 500 mM KCl (buffer B). The cells were frozen and stored at -70 degrees Celsius until protein purification.

For purification, cell pellets were thawed and mixed with two cOmplete ETDA-free protease inhibitor cocktail tablets (Roche) and 50 μL of benzonase (Molecular Biology

Services, IMP). Cells were mechanically lysed with a glass douncer, and debris was separated by centrifugation at 18,000 rpm for 30 minutes. The soluble fraction was applied to a 5 mL HP StrepTrap column (Cytiva) and washed with buffer A (25 mM HEPES pH 7.5, 150 mM KCl, 0.25 mM TCEP) and buffer B (25 mM HEPES pH 7.5, 500 mM KCl, 0.25 mM TCEP), before eluting bound protein with buffer A containing 2.5 mM d-desthiobiotin. The sample was directly loaded onto a buffer A equilibrated 6 mL ResourceQ column (Cytiva) and eluted by gradient elution towards buffer B over 20 column volumes, with BIRC6 eluting at 35-38 mSv. Protein was pooled and concentrated with a Vivaspin 30 kDa molecular weight cut-off centricon (Sartorius) and loaded onto a buffer A equilibrated S6 Increase 3.2/300 (Cytiva) via a 1 mL loop. Protein concentration was determined by 280 nm absorbance and sample aliquots were collected at every step for SDS-PAGE analysis.

### Protein production in the bacterial cells

SUMO-(MI)HTRA2 (134-358 S/A), SUMO-SMAC, XIAP or GST-LC3B plasmids were transformed into BL21 DE3 bacteria and grown in 50 mL LB media and antibiotic overnight. For expression, 2 L of LB media with either ampicillin or kanamycin was inoculated with 10 mL of the respective overnight culture, and grown to an optical density of 0.8, before inducing expression by adding 100 μM isopropyl-ß-d-thiogalactopyranoside (IPTG). Cells were grown for three hours at 37 degrees Celsius before harvesting cell pellets by centrifugation and flash frozen in liquid nitrogen.

For purifications, pellets were resuspended to a total volume of 100 mL in buffer C (50 mM HEPES pH 7.5, 300 mM KCl, 0.25 mM TCEP) and mixed with 1 tablet cOmplete ETDA-free protease inhibitor cocktail (Roche), 1 mM lysozyme, and 800 μL DNAse I (Molecular Biology Services, IMP). Proteins were loaded onto a 5 mL HisTrap (Cytiva) and washed with 20 and 50 mM imidazole in buffer C. Proteins were eluted at 300 mM imidazole in buffer C, and peak fractions pooled. SUMO constructs were cleaved overnight in dialysis. Cleaved product was re-run over the HisTrap to capture His-tagged SUMO protease, and isolate the desired product in the flow-through. Finally, samples were run on a Superose 200 16/60 (Cytiva). Fractions were again pooled, assessed for quality via SDS-PAGE, aliquoted and flash frozen in liquid nitrogen.

### Mass Photometry

Experiments were performed on a OneMP mass photometer (Refeyn Ltd.) over a one-minute acquisition time in the medium field of view via the DiscoveryMP application. Histograms of particle events were automatically Gaussian fitted with the AnalysisMP software to obtain average species masses. Samples were prepared in a 200 nM stock solution in 25 mM HEPES pH 7.5, 150 mM KCl, 0.5 mM TCEP and diluted 10x prior to measurement. Data were exported and plotted in GraphPad Prism 8 (Dotmatics).

### Ubiquitination assays

Generally, ubiquitination assays contained 9 μM wild-type ubiquitin, 1 μM DyLight488 labeled ubiquitin for in-gel visualization, 0.1 μM UBA6, 0.2 μM BIRC6 and 4 μM substrate protein, unless indicated otherwise. For LCL161 titrations, all reactions had 2% v/v DMSO. For SMAC inhibition (**Fig. 3a,d**), substrates were added at 4 μM, and SMAC was added at 0.5 μM. The LC3B∼Ub band was quantified in the presence and absence of SMAC and a ratio of activity was calculated for each respective BIRC6 mutant. The ratio was subtracted from 1 to give the relative inhibition. This was replicated four times. For SMAC ubiquitination assays (**Fig. 3e**) SMAC was added at 10 μM, and SMAC ubiquitination of BIRC6 mutants was normalized to the SMAC ubiquitination of BIRC6 WT (set to 1.0) of the individual experimental repeat. The assays were performed in 25 mM HEPES pH 7.5, 150 mM KCl, 5 mM MgCl2, 0.5 mM TCEP (assay buffer) at 37 degrees Celsius for the indicated times in 10 μL volumes. Reactions were initiated by the addition of 5 mM ATP and stopped by adding 2x reducing SDS-PAGE loading buffer. SDS-PAGE was performed using 4–12% NuPAGE Bis-Tris gradient gels (Invitrogen) in MES running buffer. Fluorescence and Coomassie imaging was performed using a Bio-Rad ChemiDoc MP system.

### Single-particle cryo-EM sample preparation

Recombinant BIRC6 was verified for homogeneity by SDS-PAGE and mass photometry. BIRC6/HTRA2 and BIRC6/SMAC complexes were prepared similarly. In 4.4 mL open-top polyclear tubes (Seton Scientific) a 10-30% sucrose gradient in 25 mM HEPES pH 7.5, 150 mM KCl, 0.5 mM TCEP was prepared with the addition of 0.15 % glutaraldehyde (Thermo Fisher Scientific) in the heavy sucrose fraction prior to gradient mixing. 50 μL of sample containing 2 μM BIRC6 and 20-30 μM ligand was added to the top of the gradient tube. Using an SW60Ti rotor in an Optima L-90K ultracentrifuge (Beckman Coulter), gradients were spun under vacuum at 26,000 rpm and 4 degrees Celsius for 16 hours before quenching with aspartate monohydrate and aliquoting 200 μL fractions. Protein presence was confirmed by amido black staining of 2 μL fraction aliquots blotted onto nitrocellulose paper (Whatman). Peak fractions were pooled and concentrated, and sucrose was removed by buffer exchanging into 25 mM HEPES pH 7.5, 150 mM KCl, 0.5 mM TCEP using Zeba Spin Desalting Columns (Thermo Fisher Scientific). Samples generated in this way were then spun down for 10 minutes at 21,000 rpm, and supernatant was transferred onto ice until further use. Protein samples were then diluted to 0.2 mg/mL shortly prior to preparation of cryo-EM grids. R3.5/1 Cu 200 grids (Quantifoil) were glow-discharged on a glass clover slide for 60 seconds at 25 mA and plunging into liquid ethane was performed using an EM GP (Leica Microsystems). 4 μL sample was applied with plunger set to 4 degrees Celsius and 75% chamber temperature and humidity, and Z-, H-and post-sensor offsets set to 3.6, 182 and +6, respectively, with a 30 second pre-blot incubation and a total blotting time of 0.22 seconds.

### Cryo-EM data collection and processing

We collected datasets for BIRC6 without a substrate, and in the presence of HTRA2, caspase-9, LC3B or SMAC. BIRC6 without a substrate was collected at the Institute of Science and Technology, Austria, using a Titan Krios G3i electron microscope (Thermo Fisher) equipped with a K3 detector (Gatan), while all other datasets were collected at the IMP using a Titan Krios G4 equipped with a Falcon 4EC detector (Thermo Fisher) using parameters described in Extended Data Table 1. The grid quality varied widely between these datasets and so to achieve a BIRC6 structure with the best overall resolution, we combined the datasets from the best quality grids, namely BIRC6:HTRA2, BIRC6:caspase-9 and BIRC6:LC3B, and refined this using C2 symmetry. We separately processed the BIRC6 dataset with no added substrate and no imposed symmetry to compare with the high resolution map and clearly distinguish those features which arise from BIRC6 from those which arise from substrates. HTRA2 was the only substrate from these three which resulted in clear additional density compared to the no substrate map, but this is mostly diluted out of the combined C2 symmetry map.

#### General approach

For the no substrate and BIRC6:SMAC complex datasets, movies were imported and then motion corrected in Relion 4.0 beta^41^, which was then used for all further processing steps. For the remaining datasets, motion correction was performed on-the-fly using Cryosparc Live^42^, and corrected images were imported into Relion 4.0 beta for all subsequent processing steps. Contrast transfer function (CTF) parameters for all datasets were determined using CTFFIND-4.1.13^43^. Manual picking and 2D classification were used to generate templates for autopicking. In general, autopicking was performed with a low threshold, and 2D classification was only used to remove ice and edges, resulting in a large number of low quality particles which were primarily removed by one or two rounds of 3D classification with alignment. Once a high-quality particle set had been obtained, conformational heterogeneity was reduced by a masked 3D classification without alignment performed with a range of T values until a class was obtained which had both the best resolution and alignment accuracy and this was selected for final 3D refinement. Due to the symmetry mismatch between BIRC6 and substrates and substrate flexibility, we found that multiple rounds of 3D classification with alignment worked better than focused classification approaches to obtain complete substrate density. Specific details for each dataset are described below.

#### BIRC6

The BIRC6:HTRA2, BIRC6:caspase-9 and BIRC6:LC3B datasets were processed separately with the same workflow. After autopicking, particles were extracted at a box size of 364 pixels (1.17 Å/pixel) and downscaled to 90 pixels (4.73 Å/pixel). The particle sets were cleaned by two rounds of 2D classification followed by one round of 3D classification with alignment, and the resulting particles were re-extracted at original pixel size. All further processing was performed with C2 symmetry imposed. For each dataset, 3D refinement was performed followed by masked 3D classification without alignment. The best classes were chosen from each dataset and combined, resulting in 240,276 particles. The same classification procedure was then repeated to yield 136,799 particles. After CTF refinement, a final resolution of 3.3 Å was achieved. Due to the large difference in local resolution across the structure, a locally sharpened map was generated, as presented in **Fig. 1**, using Locscale^44^.

#### BIRC6 – no substrate

Autopicking found 3,235,620 particles, which were extracted at a box size of 400 pixels (1.07 Å/pixel) and downscaled to 100 pixels (4.28 Å/pixel). The best classes from 2D classification were used to generate initial models with C1 and C2 symmetry. For further processing only ice, edge and carbon classes were removed using 2D classification to leave 1,932,830 particles. One round of 3D classification with alignment and C1 symmetry removed further poor quality particles leaving 1,091,933 particles. A second round of 3D classification with alignment with imposed C2 symmetry was used to remove further particles without the distinctive BIRC6 shape resulting in 258,934 particles. Particles were reextracted without binning and 3D refinement and particle polishing were then performed in C2 symmetry. C1 symmetry was used for the final processing steps. First, 3D refinement was performed followed by a 3D classification without alignment. The best class was refined with a mask to a resolution of 7.0 Å, judging by an FSC of 0.143.

#### BIRC6:HTRA2 complex

Autopicking from 8,842 movies resulted in 1,879,003 particles which were extracted at a box size of 364 pixels (1.17 Å/pixel) and downscaled to 90 pixels (4.73 Å/pixel). 1,152,643 protein particles remained after 2D classification, which were further cleaned by one round of unmasked 3D classification with alignment leaving 226,837 particles with the distinctive BIRC6 shape which were reextracted at their original pixel size. Extra density in the central pore was already clear at this stage. Two rounds of 3D classification with alignment were performed and classes where the bound density had three-fold symmetry were selected, leaving 15,780 particles. Masked 3D refinement yielded a map with a final resolution of 6.2 Å. This was low-pass filtered to either 8 or 10 Å to better visualize the HTRA2 homotrimer.

#### BIRC6:SMAC complex

Three separate data collections from the same grid were processed separately until the first round of 3D classification after which they were combined. Otherwise, the processing pipeline was approximately the same as for the BIRC6:HTRA2 complex. Autopicking found a total of 8,724,828 particles which were extracted at a box size of 574 pixels (0.7445 Å/pixel) and downscaled to 100 pixels (4.27 Å/pixel). After the first round of 3D classification, particles were re-extracted with a 300 pixel box size (1.42 Å/pixel) and the datasets combined for a total of 599,274 particles. Classes with poor SMAC or BIRC6 density were removed through two further rounds of 3D classification with alignment, leaving 168,962 particles. 3D refinement was then performed prior to focused classification using a mask a soft mask encompassing SMAC and the nearby regions from BIRC6. One class showed high resolution SMAC features and was selected for final masked 3D refinement yielding a map at 8.4 Å resolution.

### Model building

The BIRC6 structure was modelled and refined using the high resolution C2 symmetry map with either *B* factor blurring (100 or 500 Å^2^), for the lower resolution exterior regions or *B* factor sharpening (−27.1 Å^2^) for the higher resolution core. Initially, three overlapping models were generated using a locally run version of ColabFold^45,46^ for the residue ranges 1-2280, 1001-3229 and 2221-4500. These could be directly docked into the experimental density using UCSF ChimeraX^47^ with some rigid body fitting of local sections using Coot^48^ due to the long-scale flexibility of the Armadillo repeats. However, regions that formed the dimer interface were incorrect. Once this interface had been identified, a homodimer model was calculated using the input sequence of residues 2090-3447 without the sections between 2192-2321 and 2422-2631. This fit very well to the experimental density. Further rounds of building and refinement were performed using Coot and Phenix Real Space Refine^49^.

For the BIRC6 no substrate structure, the above model was docked into the unsharpened density using UCSF ChimeraX, followed by rigid body fitting of local sections in Coot and refinement using Phenix Real Space Refine.

The BIRC6:HTRA2 map was blurred with a *B* factor of 200 Å^2^. The above BIRC6 model and the crystal structure of the HTRA2 (S306A) trimer without PDZ domains (PDB:7VGE) were then docked into the density using UCSF ChimeraX, followed by rigid body fitting of local sections of BIRC6 in Coot. The model was then refined using Phenix Real Space Refine.

The BIRC6:SMAC map was sharpened with a *B* factor of -100 Å^2^. BIRC6 was modelled and refined with the same procedure as the BIRC6 no substrate structure.

A Colabfold model of dimeric SMAC was docked in the additional density. The N-degron was built using PDB:1G73. The CC1 insert (2226-2303) identified by XL-MS was fit into the remaining density using the 1001-3229 Colabfold sequence block, taking the BIRC6 chain where the insert had the shortest physical distance from where it emerges in the Armadillo backbone. Local rigid body fitting was performed using Coot followed by global refinement with Phenix Real Space Refine.

### Negative stain EM

Protein samples were diluted to 0.08 mg/mL and continuous carbon grids glow discharged at 25 mA for 60 seconds using a BAL-TEC sputter coater CD005 on a glass cover slide. 4 uL of protein was added and incubated for 60 seconds, blotted to near dryness with filter paper (Whatman), washed with 4 μL 2% uranyl acetate, blotted again, and incubated for 60 seconds with 4 μL 2% uranyl acetate before blotting to complete dryness. Grids were examined on an 80 kV FEI Morgagni and a 200 kV FEI Tecnai G2 20 (Thermo Fisher Scientific). For initial 2D and 3D classes of BIRC6, a T20 dataset at x62,000 magnification at a pixel size of 1.85 Å was collected which yielded 36,000 particles for downstream processing in Relion 3.1.

### Cross-Linking Mass-Spectrometry

Sample preparation: 2 μM of BIRC6 alone or with equimolar amounts of SMAC was prepared in 25 mM HEPES pH 8, 50 mM KCl. The protein sample was crosslinked with 50 μM BS3 (bis(sulfosuccinimidyl)suberate) (Thermo Fisher Scientific) for 30 min at 37 degrees Celsius and 500 rpm. The reaction was quenched by adding ammonium bicarbonate to a concentration of 50 mM and incubating the mixture for 10 min at 37 degrees Celsius and 500 rpm. The sample was vacuum dried at 45 degrees Celsius. and resuspended in aqueous 8 M urea. The mixture was reduced with 2.5 mM TCEP for 30 min at 37 degrees Celsius and 850 rpm. Next, the proteins were alkylated with 5 mM iodoacetamide and incubated for 30 min at room temperature in the dark. After digesting the sample with 1.2 μg LysC per 100 μg protein for 4 h at room temperature, urea was diluted to 1 M by the addition of 50 mM ammonium bicarbonate. The proteins were then further digested with 2 μg MS-grade trypsin (Thermo Fisher Scientific) per 100 μg protein for 20 hours at 37 degrees Celsius and 850 rpm. After this, trypsin was deactivated with 1 % TFA. The peptide fragments were then loaded onto a Sep-Pak C18 cartridge (Waters) equilibrated with 5% acetonitrile, 95% water, 0.1% formic acid. The sample was eluted with 50% acetonitrile, 50% water, 0.1% formic acid. For additional purification by size exclusion chromatography, the peptides were dried at 45 degrees Celsius, resuspended in 30% acetonitrile, 70% water, 0.1 % triformic acid and loaded onto a Superdex 30 Increase 3.2/300 column (Cytiva). Fractions were pooled, dried at 45 degrees Celsius. and stored at -20 degrees Celsius until analysis.

LC-MS/MS: Generated peptides were injected into a PepMap C18 5 mm x 300 um column (Thermo Fisher Scientific) at a flow of 25 μL/min with 0.1% TFA mobile phase. After 10 minutes flow was switched toward an analytical column PepMap C18, 500 mm x 75 uM (Thermo Fisher Scientific) operated at a flow of 230 nL/min at 30 degrees Celsius. Peptides were separated under a solvent gradient between buffer A and B (0.1% formic acid in water and 80% acetonitrile, 0.08% formic acid in water, respectively), starting at 2% buffer B, before increasing to 35% over 60 minutes and peaking at 90% for 5 minutes, before decreasing to 2% again. Peptides were analyzed on an Ultimate 3000 RSLCnano HPLC, coupled by a Nanospray Flex ion source to an FAIMS pro interface equipped Obritrap Exploris 480 mass spectrometer (Thermo Fisher Scientific). Operated in data-dependent mode, the Orbitrap performed a full scan (375-1500 m/z range at 60,000 resolution and target value 3E6) and CVs of -40, -50, -60, with a resulting MS/MS scan cycle time of 1 second/CV. Spectra were acquired with a stepped normalized collision energy of 28+/-2%, 1.4 m/z isolation width and 30,000 resolution. The fill time of 100 miliseconds and intensity threshold of 1E4 and a fixed first mass of 120 = m/z was used. Exclude isotope features was set to enabled and the peptide match feature was set to preferred.

pLink 2.3.9 was used for analysis of raw files. MS/MS spectra were queried against a database containing BIRC6 and SMAC, along with a pLink derived common contaminant list. A fragment and peptide mass tolerance was set to ± 10 ppm. The minimum peptide length was set to 6 residues with a maximum mass of 1000 Dalton. BS3 was defined as a lysine cross-linker to also include monolinks. Variable modification was set for glutamines and asparagines to cover deamidation, methionine oxidation, and cysteine carbamidomethylation. Data was filtered to FDR 1% by pLink and the CSM E value to <0.001. Hits were visualized as 2D network in web-based Xiview.

### Statistics and reproducibility

The experiments shown in **Figs. 1b,c,d**, **2c,d,e** and **3a** were independently performed at least two times. For each replicate the same result was result achieved and no data were excluded. For experiments in **Fig. 3d** and **3e**, the data from all independently performed replicates are shown.

### Reporting Summary

Further information on research design is available in the Nature Research Reporting Summary linked to this article.

## Data availability

Cryo-EM maps and atomic coordinates have been deposited in the Protein Data Bank (PDB) with accession codes 8ATU, 8ATX, 8AUK and 8AUW and the Electron Microscopy Data Bank (EMDB) with accession codes: EMD-15663, EMD-15668, EMD-15672, EMD-15675.

## EXTENDED DATA

**Extended Data Fig 1.**
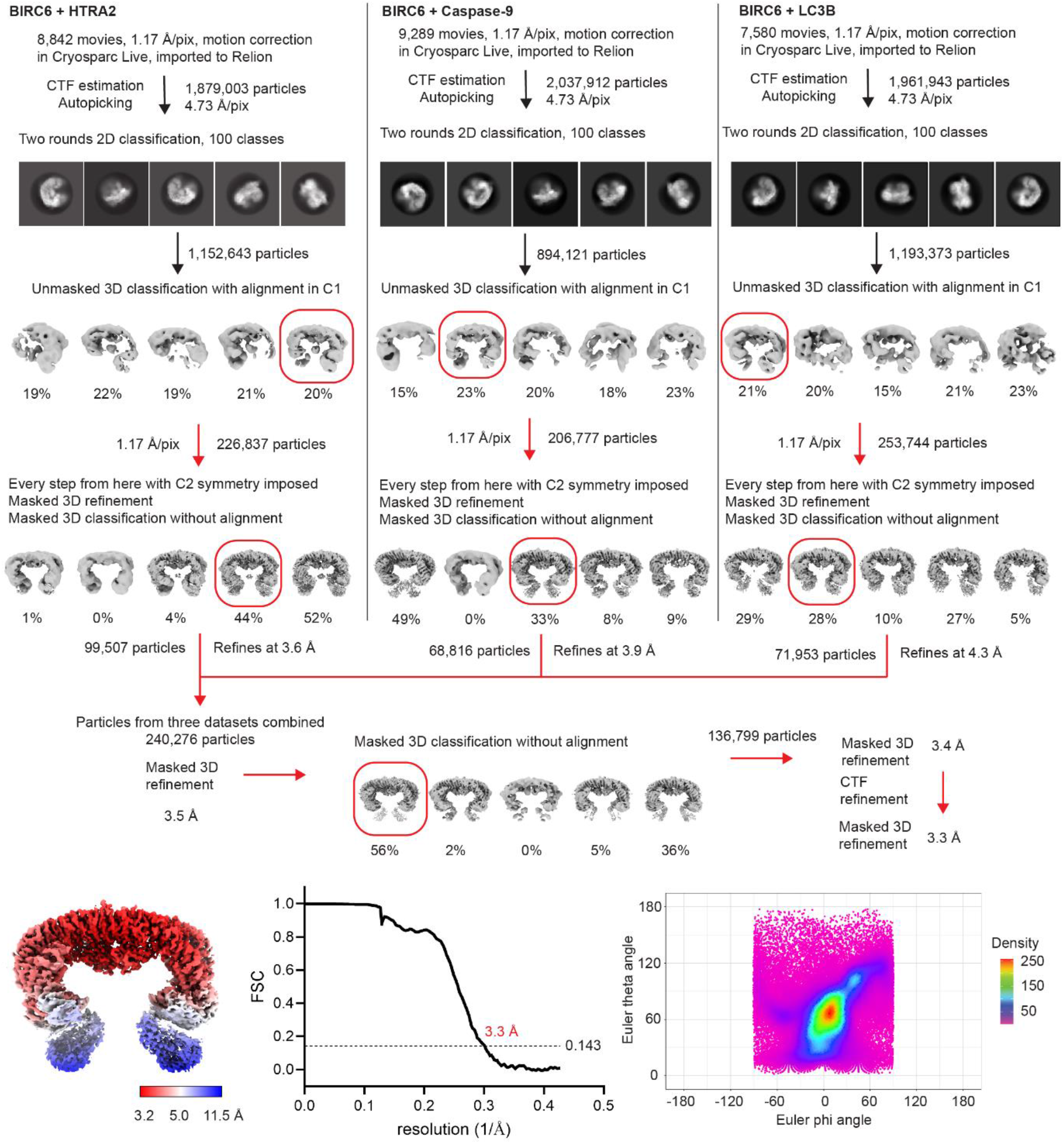
Processing workflow to obtain a high resolution BIRC6 structure. Details are provided in the Methods section. In the lower panels, the local resolution, masked and corrected FSC curve and angular distribution of alignments of the final model are shown.

**Extended Data Fig 2.**
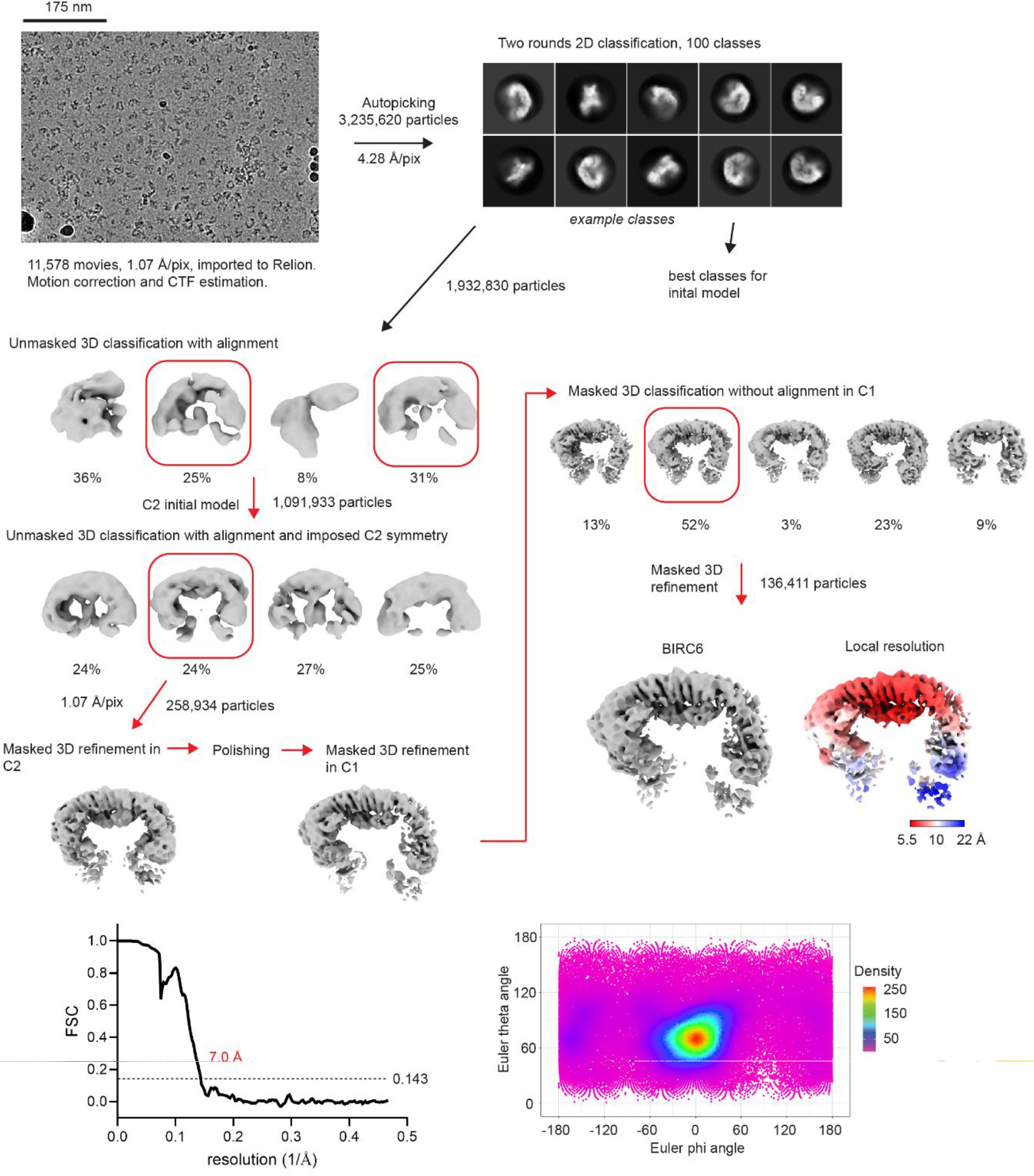
Processing workflow for BIRC6 in the absence of substrates. All details are provided in the Methods section. In the lower panels, masked and corrected FSC curve and angular distribution of alignments for the final model are shown.

**Extended Data Fig 3.**
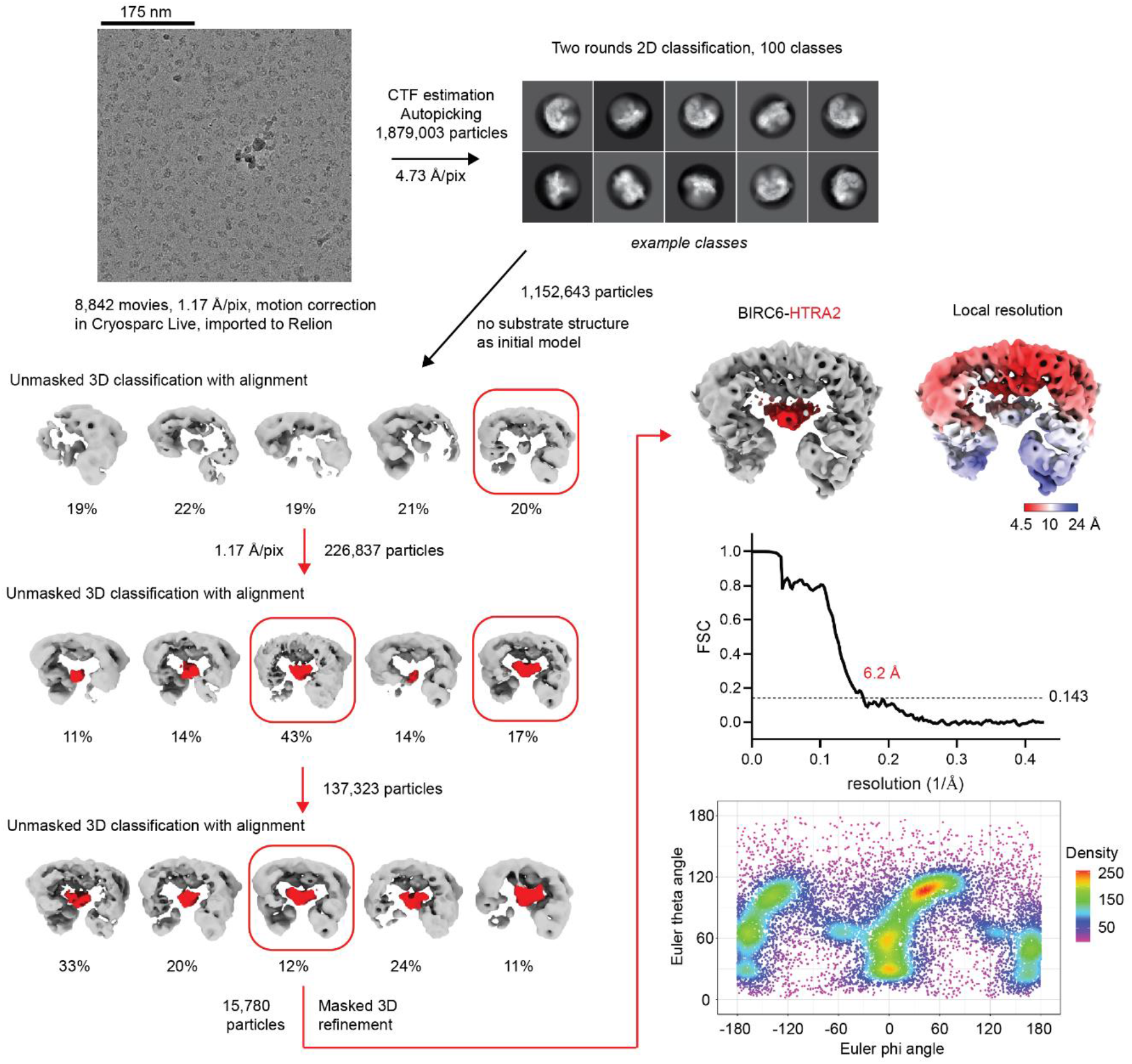
Processing workflow for the BIRC6:HTRA2 complex dataset. Details are provided in the Methods section. In the last panels, the final masked and corrected FSC curve and angular distribution contributing to the final model are shown.

**Extended Data Fig 4.**
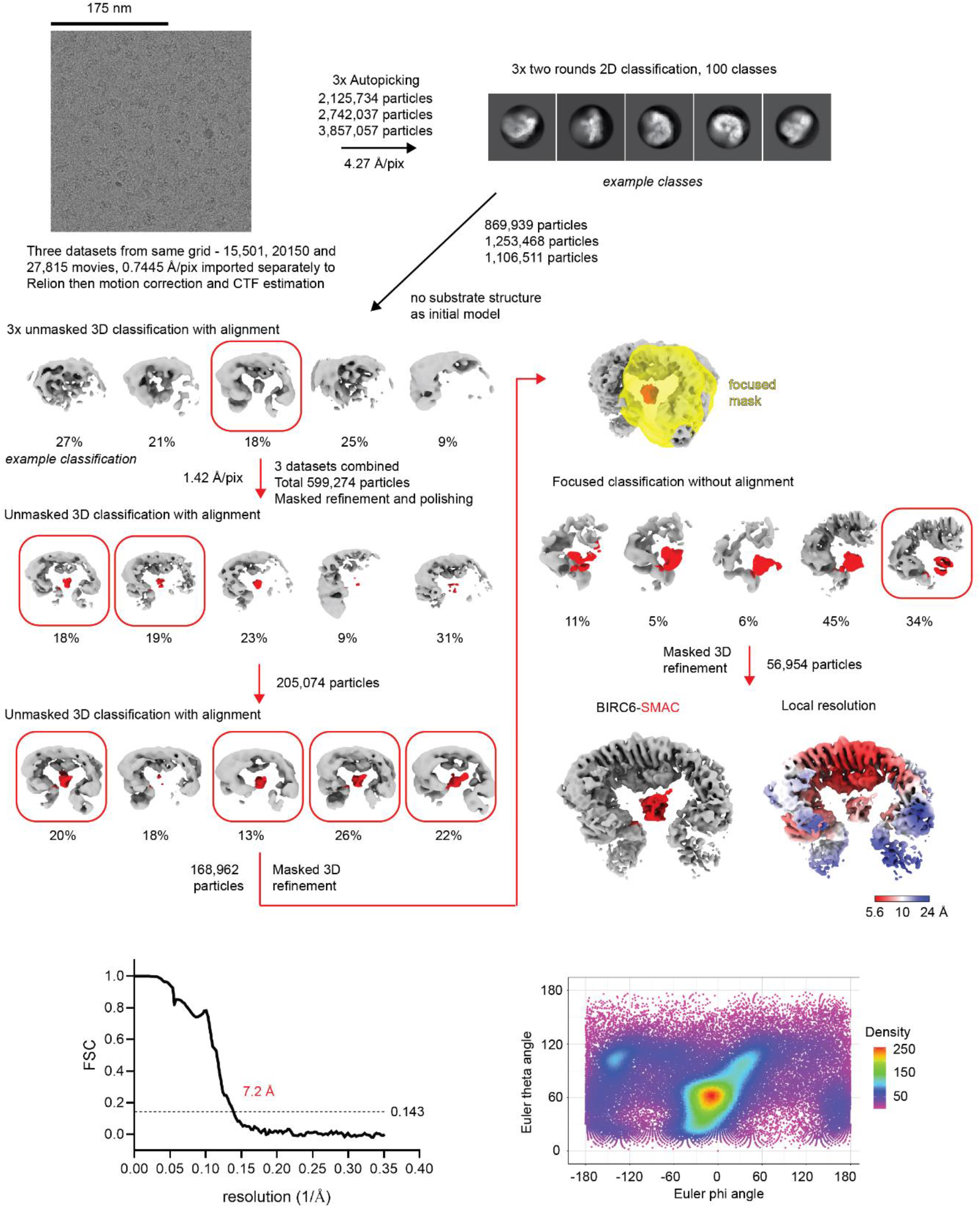
Processing workflow for the BIRC6:SMAC complex dataset. All details are provided in the Methods section. In the lower panels, the masked and corrected FSC curve and angular distribution of the final model are shown.

**Extended data Fig 5.**
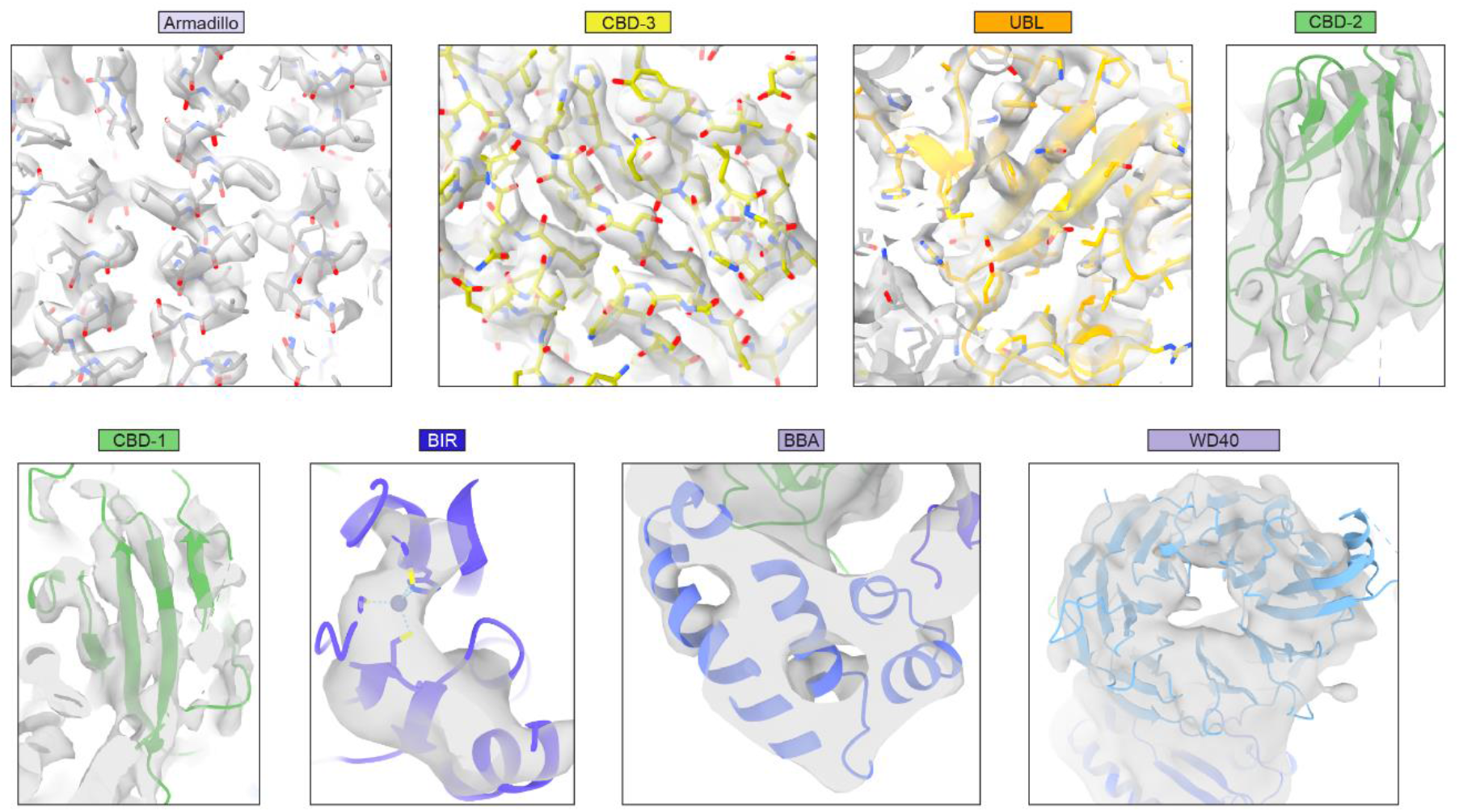
Fit of the model to the density in the different regions of the map. Maps with different *B* factor sharpening values are shown due to the wide range of local resolution. The N-terminal module was best resolved in the BIRC6:SMAC complex map. The specific maps shown are as follows: Armadillo and CBD3 – BIRC6 main map (*B* -27.1 Å^2^), UBL – BIRC6 main map (*B* 0 Å^2^), CBD2 and CBD1 – BIRC6 main map (*B* +100 Å^2^), BIR and BBA – BIRC6:SMAC complex (*B* -100 Å^2^), WD40 – BIRC6:SMAC complex (*B* +100 Å^2^).

**Extended Data Fig 6.**
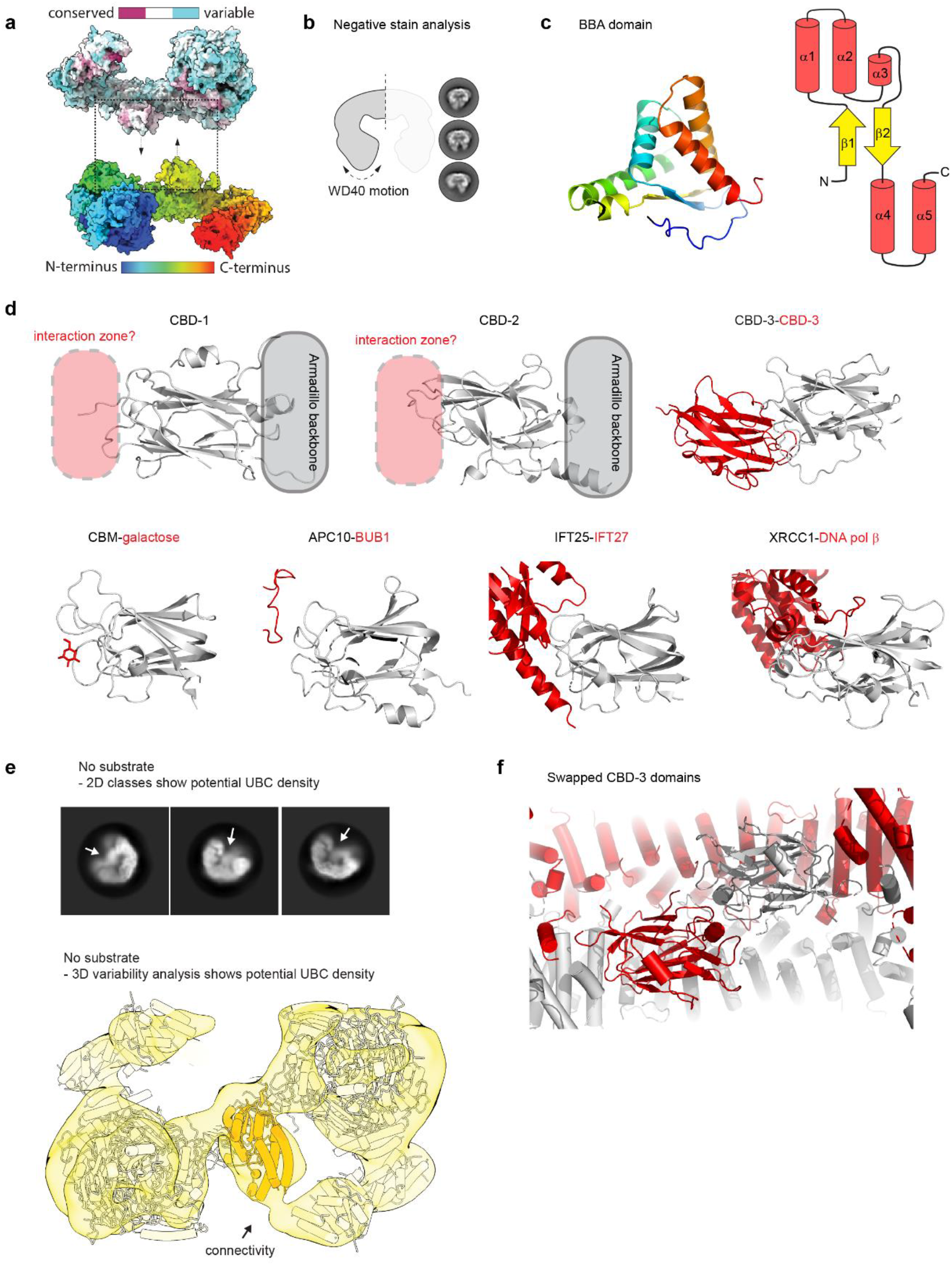
Structural features of BIRC6. **a**, The two BIRC6 protomers are separated and shown in surface representation. The upper is colored by sequence conservation, while the lower is shown in rainbow coloring from N-to C-terminus. **b**, Negative stain analysis of BIRC6 shows flexibility of the N-terminal modules. Aligned 2D classes are shown. **c**, Fold and topology of the novel BBA domain. The structure is shown in rainbow coloring on the left from N-terminus (blue) to C-terminus (red), and using a topology diagram on the right. **d**, Structural comparison of the BIRC6 CBD domains with structural homologues identified using the DALI server^50^. Other CBD domains interact using a similar surface with sugars, peptides and proteins, that in CBD-1 and CBD-2 is pointing into the central cavity, and for CBD-3 is used for homodimerization. The CBD is always shown in grey, and the interactor in red. The lower panel shows from left to right: PDB 7BLG, PDB 5LCW, PDB 2YC2 and PDB 3K75. **e**, The flexible UBC domains may enter the central BIRC6 cavity. Both panels show analysis of the BIRC6 dataset collected in the absence of substrates. Many 2D classes have heterogeneous density in the central area of BIRC6 which is likely to be the UBC domain as this is the only unmodelled domain. Cryosparc 3D variability analysis reveals continuous density with the approximate dimensions of the UBC domain connected to the C-terminus of the armadillo repeat. **f**, The CBD-3 domains are swapped across BIRC6 protomers, stabilizing the dimer. The two BIRC6 protomers are shown in grey and red.

**Extended Data Fig 7.**
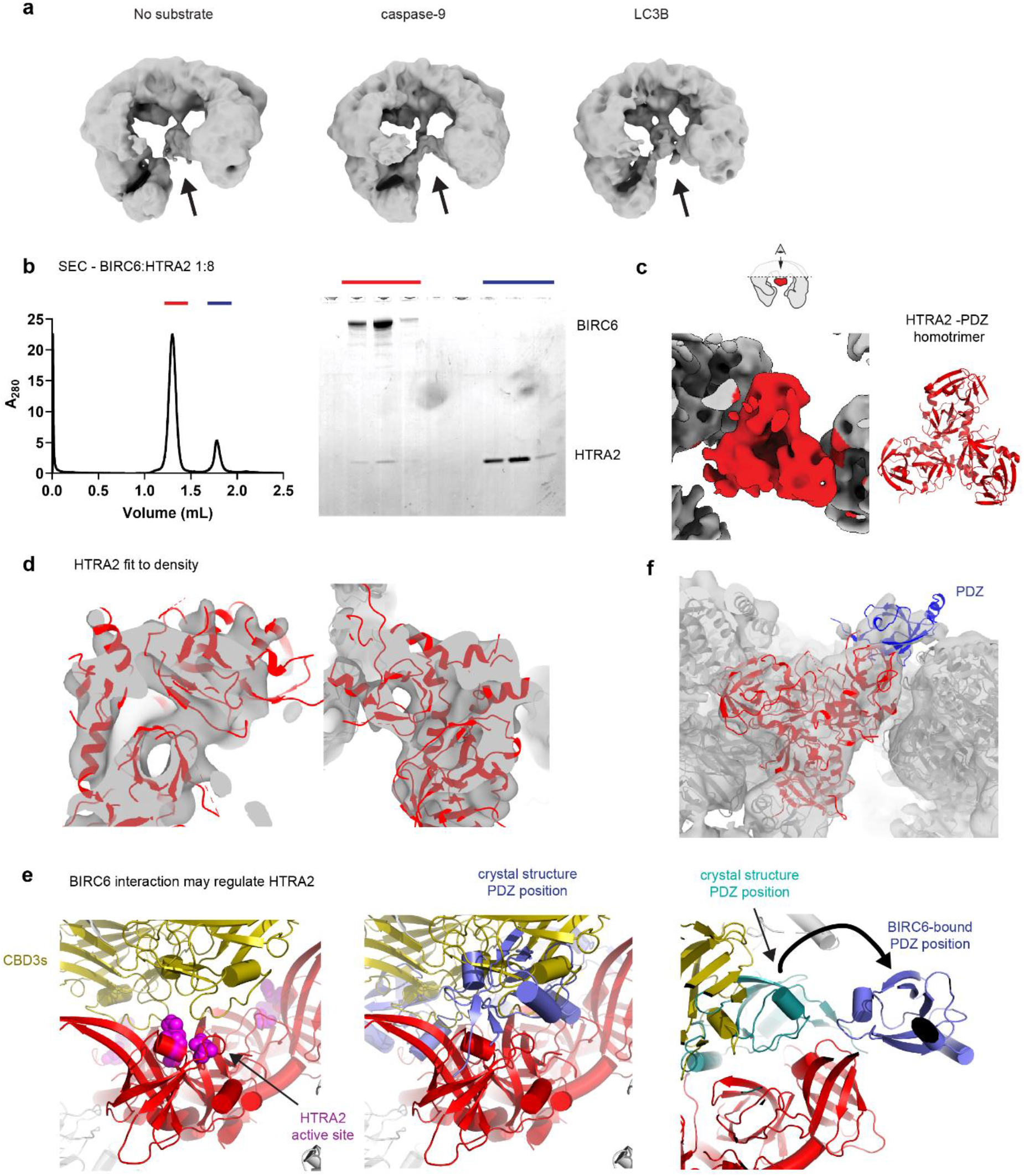
Features of the BIRC6:HTRA2 complex structure. **a**, Maps of BIRC6 in the absence of substrates or with caspase-9 or LC3B all show heterogeneous density in the central cavity. In each case, the map is the first refinement using a spherical mask after initial filtering of poor quality particles. **b**, BIRC6 and HTRA2 form a stable complex in size-exclusion chromatography. Fractions corresponding to the two peaks were analyzed by SDS-PAGE and show co-elution in the main peak. **c**, A cross-section of the BIRC6:HTRA2 map lowpass filtered to 10 Å resolution reveals the characteristic shape of the HTRA2 homotrimer, with the cartoon model of HTRA2 for comparison on the right. **d**, Local features of HTRA2 such as β-barrels and some helices can also be observed in the density lowpass filtered to 8 Å resolution, confirming the fit. **e**, How the CBD-3-HTRA2 interaction may regulate HTRA2. In the left panel the HTRA2 catalytic triad is highlighted in sphere representation showing that it faces CBD-3. In the middle panel, the crystal structure of full-length HTRA2 (PDB 1LCY) is overlaid on our cryo-EM structure, revealing a clash between the position of PDZ domains in the resting state with the CBD-3 domains of BIRC6. **f**, Density consistent with a displaced PDZ is visible in our BIRC6:HTRA2 map, which is shown lowpass filtered to 10 Å resolution. A cartoon shows how the PDZ domain would have to rotate 180° out of its resting state position in PDB 1LCY to reach the position we observe.

**Extended data Fig 8.**
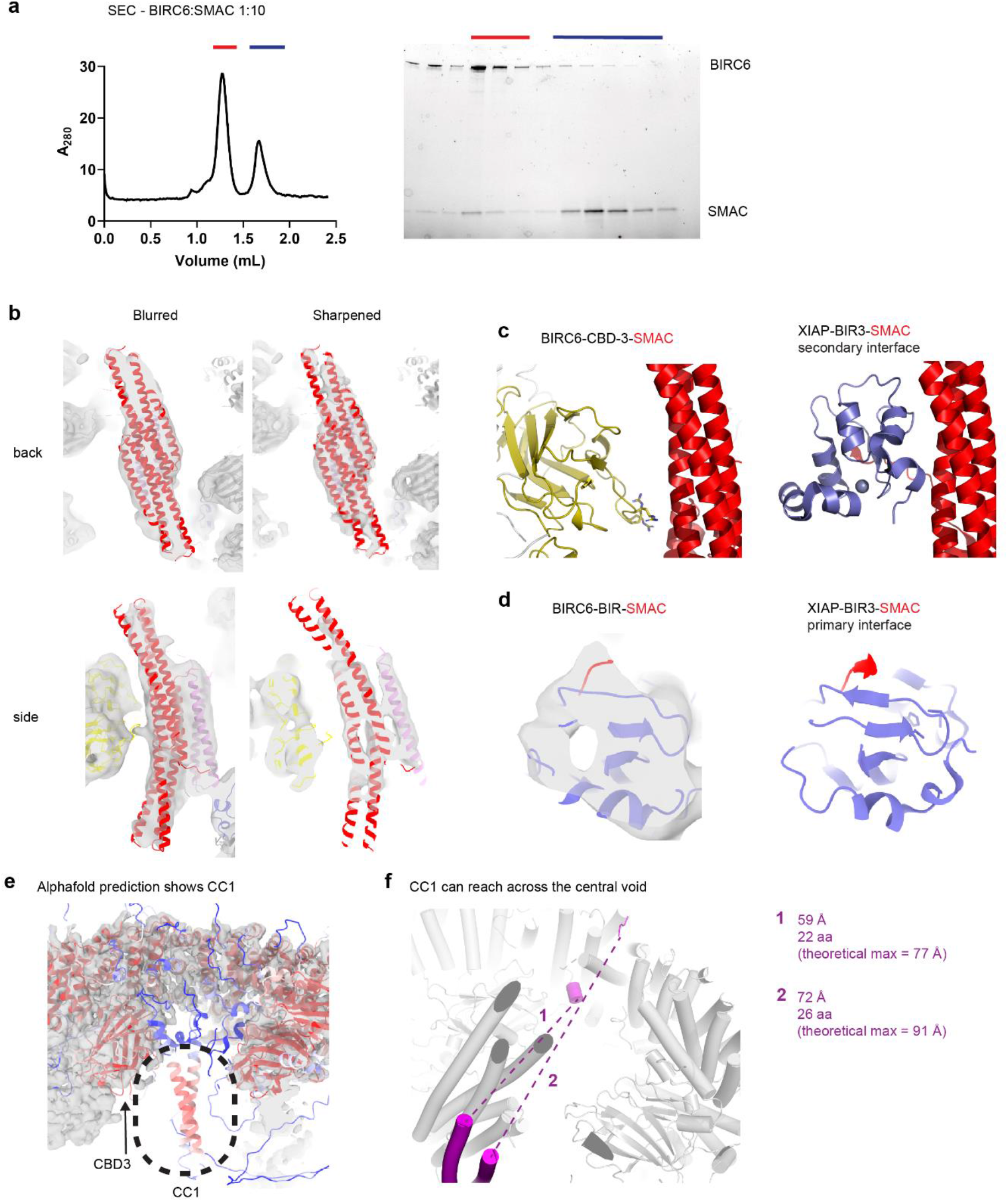
Features of the BIRC6:SMAC complex structure. **a**, BIRC6 and SMAC form a stable complex in size-exclusion chromatography. SDS-PAGE shows co-elution of the two proteins in the first peak. **b**, The SMAC homodimer can be unambiguously placed in the extra density. The density is shown either *B* factor sharpened (−100 Å^2^) to highlight higher resolution features or blurred (+100 Å^2^) to show that the overall shape of the additional density is consistent with a SMAC dimer. **c** and **d**, comparison of the SMAC binding modes in our BIRC6:SMAC complex structure with the previous structure of the XIAP BIR3 domain in complex with SMAC (PDB 1G73). The *B* factor sharpened (−100 Å^2^) BIRC6:SMAC map is shown in the left panel of **d. e**, An Alphafold prediction of BIRC6 residues 1001-3229 shows a confidently predicted coiled coil in central cavity of BIRC6. The model is colored by pLDDT score and shown docked in the main BIRC6 map. **f**, The modeled position of CC1 is shown in relation to where this insert emerges from the armadillo backbone. The theoretical maximum distance that an extended stretch of residues is estimated at 3.5x the number of amino acids.

**Extended data Table 1.** Data collection and refinement statistics.

|  | BIRC6<br>PDB 8ATU,<br>EMD-15663 | BIRC6 – no<br>substrate<br>PDB 8ATX,<br>EDM15668 | BIRC6:HTRA2<br>PDB 8AUK,<br>EMD-15672 | BIRC6:SMAC<br>PDB 8AUW,<br>EMD-15675 |
| --- | --- | --- | --- | --- |
| **Data collection and processing** |  |  |  |  |
| Magnification | 105,000x | 81,000x | 105,000x | 165,000x |
| Voltage (kV) | 300 | 300 | 300 | 300 |
| Electron exposure (e<sup>-</sup><br>/Å<sup>2</sup>) | 60 | 60 | 60 | 41 |
| Defocus range (μm) | -0.5 – -3.0 | -0.9 – -3.0 | -0.5 – -3.0 | -0.5 – -1.5 |
| Pixel size (Å) | 1.17 | 1.07 | 1.17 | 1.42 |
| Symmetry imposed | C2 | C1 | C1 | C1 |
| Initial particle images<br>(no.) | 5,878,858 | 3,235,620 | 1,879,003 | 8,724,828 |
| Final particle images<br>(no.) | 136,799 | 136,411 | 15,780 | 56,954 |
| Map resolution (Å) | 3.3 | 7.0 | 6.2 | 7.2 |
| FSC threshold | 0.143 | 0.143 | 0.143 | 0.143 |
| Map resolution range | 3.2-11.5 | 5.5-22 | 4.5-24 | 5.6-24 |
| **Refinement** |  |  |  |  |
| Initial model used | AlphaFold | BIRC6 (8ATU) | BIRC6 (8ATU)<br>PDB:7VGE | BIRC6 (8ATU),<br>AlphaFold, PDB:<br>1G73 |
| Model resolution (Å) | 3.7 | 8.4 | 7.9 | 8.4 |
| FSC threshold | 0.5 | 0.5 | 0.5 | 0.5 |
| Map sharpening *B*<br>factor (Å<sup>2</sup>) | -27.1 | 0 | +200 | -100 |
| **Model composition** |  |  |  |  |
| Non-hydrogen atoms | 43,882 | 43,882 | 47,827 | 46,881 |
| Protein residues | 5,674 | 5,674 | 6204 | 6,049 |
| Ligands | 2 x Zn<sup>2+</sup> | 2 x Zn<sup>2+</sup> | 2 x Zn<sup>2+</sup> | 2 x Zn<sup>2+</sup> |
| ***B* factors (Å<sup>2</sup>)** |  |  |  |  |
| Protein | 150.6 | 438.8 | 413.6 | 542.0 |
| Ligand | 338.4 | 879.9 | 667.8 | 768.8 |
| **R.m.s. deviations** |  |  |  |  |
| Bond lengths (Å) | 0.003 | 0.007 | 0.006 | 0.006 |
| Bond angles (°) | 0.763 | 0.912 | 1.130 | 1.151 |
| **Validation** |  |  |  |  |
| MolProbity score | 1.62 | 1.71 | 1.74 | 1.61 |
| Clashscore | 7.12 | 9.58 | 11.44 | 10.20 |
| Poor rotamers (%) | 0.4 | 0.0 | 0.0 | 0.0 |
| **Ramachandran plot** |  |  |  |  |
| Favored (%) | 96.54 | 96.72 | 97.08 | 97.64 |
| Allowed (%) | 3.43 | 3.25 | 2.88 | 2.34 |
| Disallowed (%) | 0.04 | 0.04 | 0.03 | 0.02 |

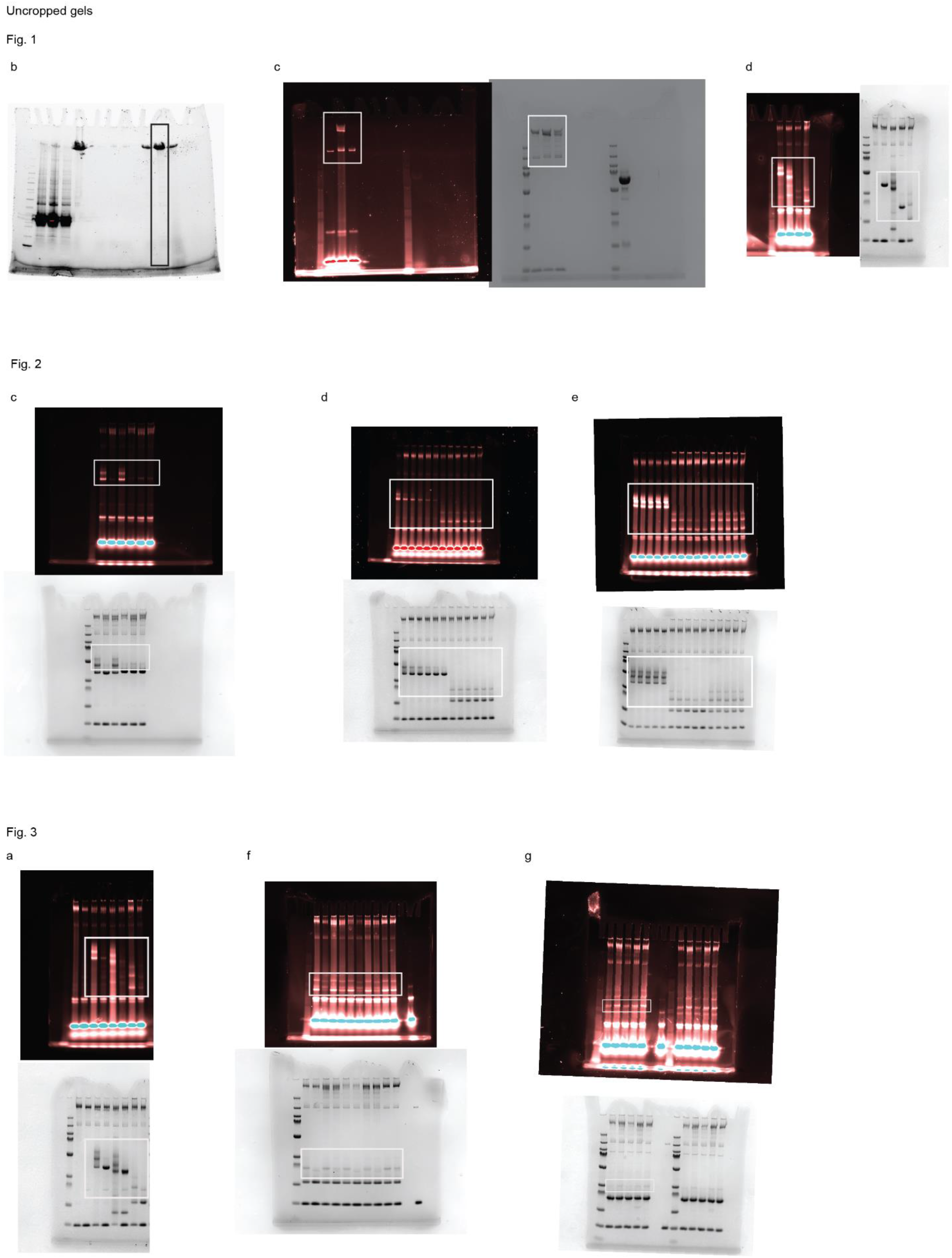

